# Synthesis of oxyfunctionalized NSAID metabolites by microbial biocatalysts

**DOI:** 10.1101/315374

**Authors:** Jan M. Klenk, Lisa Kontny, Bernd A. Nebel, Bernhard Hauer

**Affiliations:** Institute of Biochemistry and Technical Biochemistry, University of Stuttgart, Allmandring 31, 70569 Stuttgart, Germany

**Keywords:** Biotransformation, Filamentous fungi, Microbial biocatalysts, NSAIDs, Oxyfunctionalized metabolites

## Abstract

The synthesis of valuable metabolites and degradation intermediates of drugs, like non-steroidal anti-inflammatory drugs (NSAIDs), are substantially for toxicological and environmental studies, but efficient synthesis strategies and the metabolite availability are still challenging aspects. To overcome these bottlenecks filamentous fungi as microbial biocatalysts were applied. Different NSAIDs like diclofenac, ibuprofen, naproxen and mefenamic acid could be oxyfunctionalized to produce human metabolites in isolated yields of up to 99% using 1 g L^−1^ of substrate. Thereby the biotransformations using *Beauveria bassiana*, *Clitocybe nebularis* or *Mucor hiemalis* surpass previous reported chemical, microbial and P450-based routes in terms of efficiency. In addition to different hydroxylated compounds of diclofenac, a novel metabolite, 3’,4’-dihydroxydiclofenac, has been catalyzed by *B. bassiana* and the responsible P450s were identified by proteome analysis. The applied filamentous fungi present an interesting alternative, microbial biocatalysts platform for the production of valuable oxyfunctionalized drug metabolites.

**Importance:** The occurrence of pharmaceutically active compounds, such as diclofenac and its metabolites, in the environment, in particular in aquatic systems, is of increasing concern because of the increased application of drugs. Standards of putative metabolites are therefore necessary for environmental studies. Moreover, pharmaceutical research and development requires assessment of the bioavailability, toxicity and metabolic fate of potential new drugs to ensure its safety for users and the environment. Since most of the reactions in the early pharmacokinetics of drugs are oxyfunctionalizations catalysed by P450s, oxyfunctionalized metabolites are of major interest. However, to assess these metabolites chemical synthesis often suffer from multistep reactions, toxic substances, polluting conditions and achieve only low regioselectivity. Biocatalysis can contribute to this by using microbial cell factories. The significance of our research is to complement or even exceed synthetic methods for the production of oxyfunctionalized drug metabolites.

## Introduction

Next to chemical processes, biotechnology routes can play a significant role for the access to pharmaceutically active substances and their metabolites. Metabolites and degradation intermediates can have major environmental implications, which can lead to unwanted side reactions in nature and mammalians. Due to the fact that numerous drug metabolites are detected in higher amounts in different habitats, thus detailed environmental and toxicological studies are necessary. These findings induced that the FDA (Food and Drug Administration) issued guidelines for metabolites in drug testing, setting a limit of 10% in 2008.

Approximately 80% of all reactions in the early pharmacokinetics of drugs (phase I reactions) are P450-catalyzed (1, 2). Therefore, oxyfunctionalized metabolites are often major degradation intermediates of potential toxic endogenous and exogenous compounds in mammalians and nature. For further approval of new drugs, knowledge about potential metabolism intermediates and their availability for toxicological studies are obligatory (3–5). As a very present example, a molecule like diclofenac (**1**), a widespread non-steroidal anti-inflammatory drug (NSAID), and its already known hydroxylated degradation metabolites 4’- and 5-hydroxydiclofenac (**2** and **3**) pose an environmental problem due to their low degradability (6, 7). Thereby, the request for particular drug metabolites increases simultaneously with the number of newly developed drugs, to study pharmacokinetic and toxic effects caused by metabolism (8, 9). Due to limited availability and often time as well as cost elaborating chemical synthesis routes, the value of drug metabolites can reach up to several hundred dollars per milligram. As mentioned above, the chemical synthesis of human metabolites such as 4’- and 5-hydroxydiclofenac (**2** and **3**) for toxicological studies is tedious and has only been achieved in low yields linked with high by-product formation (10–12). To complement synthetic methods, microbial processes with heterologous expressed P450s and wild-type organisms have emerged as an valuable alternative to chemical syntheses (13, 14).

Most of the previous studies focused on heterologous expressed human P450s which can produce the corresponding metabolites detectable in the human body (15). One example is CYP2C9 that was recombinant produced in different hosts and finally used for the production of 468 mg L^−1^ of 4’-hydroxydiclofenac (**2**) (14, 16). Furthermore, the low activities and selectivities of the wild type P450 BM3 were improved by directed evolution resulting in variants which led to a significantly enhanced oxidation capability of NSAIDs (17–20). In this context we recently published the scale-up of the self-sufficient P450 RhF (21). The enzyme was heterologous expressed and implemented as whole cell system, whereby 5-hydroxydiclofenac (**3**) was produced exclusively in high titers of up to 357 mg L^−1^. However, high yields with NSAIDs are rather the exception, because often the product yields obtained with P450s or microorganisms are in the lower milligram range (17, 22–26). Therefore, the biotechnological synthesis of such oxyfunctionalized compounds for environmental and toxicological studies remains expandable.

Based on our previous hydroxylation experiment studies with diclofenac (**1**) as substrate, we further extended the biocatalyst platform for NSAID oxyfunctionalizations by a spectrum of available biocatalysts. In this work, we intended to identify new microbial biocatalysts able to catalyze NSAID metabolites and degradation intermediates in high yields, to provide those molecules for further drug development and toxicological studies.

## Materials and Methods

### Materials

The chemicals and media used in this study were obtained from Fluka (Buchs, Switzerland), Sigma-Aldrich (St. Louis, Missouri, USA) Alfa-Aesar (Ward Hill, Massachusetts, USA) and Carl-Roth (Karlsruhe, Germany) in highest available purity degrees. The KOD HS polymerase was from Novagene Inc. (Madison, Wisconsin, USA).

### Isolation of genomic DNA from eukaryotes

For the isolation of genomic DNA from eukaryotes, precultures of the various filamentous fungi were cultured on PEG broth (potato-extract-glucose broth) as medium for about 3 days at 27°C and 180 rpm. An initial cell lysis was achieved by centrifugation of the cultures, freezing at −80°C for 10 min and crushing the frozen mycelia with a mortar. Subsequently, the samples were isolated according to the manufacturer’s instructions using the ZR Fungal/Bacterial DNA Microprep ™ kit (Zymo Research Corp., Irvine, California, USA). The elution of the DNA was carried out with 25 μL ddH_2_O.

### Polymerase chain reaction (PCR) for the amplification of DNA fragments

The PCR was used to selectively amplify DNA fragments of genomic DNA based on the primers ITS4 (5’-TCCTCCGCTTATTGATATGC-3’) and ITS5 (5’-GGAAGTAAAAGTCGTAACAAGG-3’) or LR0R (5’-ACCCGCTGAACTTAAGC-3’) and LR5 (5’-TCCTGAGGGAAACTTCG-3’) as described by Schoch and coworkers in 2012 (27). For the elongation KOD HS Polymerase was used in 50 μL total volume using the components and programs as shown in TABLE S1 and S2.

The success of the PCR reactions was controlled by an agarose gel and the DNA fragments were then directly isolated (Figure S1). Therefore, the DNA fragments were visualized under UV light (366 nm) and cut out of the agarose gel with a scalpel, weighed and treated with the Zymoclean Gel DNA Recovery Kit (Zymo Research Corp., Irvine, California, USA) according to the manufacturer’s instructions. The isolated DNA was subsequently eluted with 20 μL ddH2O. For the analysis of the DNA samples, a volume of 20 μL and a concentration of 50 - 70 ng μL^−1^ were used. The sequencing was done by GATC Biotech AG (Konstanz, Germany) using the primers of the PCR amplification.

### Proteome analysis of Beauveria bassiana

For the cell disruption of *Beauveria bassiana*, liquid cultures were centrifuged and the mycelia subsequently washed with ddH20 over a filter paper and dried. In a Petri dish, the samples were lyophilized (Alpha 2-4 LD plus, Martin Christ Gefriertrocknungsanlagen GmbH, Osterode am Harz, Germany), to rub this later as a first partial disruption with the mortar. 35-50 mg of powder was mixed with an equivalent volume of glass beads (0.1-0.25 mm) and 1 mL urea buffer (28) (25 mM Tris/HCl pH 6.8, 9 M urea, 1% SDS, 1 mM EDTA, direct before use 0.7 M DTT). Subsequently, the cell disruption samples were heated to 95°C for 2 min, shaken for 1 min by vortexing and heated again at 95°C for 1 min. After separation of the glass beads by short centrifugation, the DNA was sheared by sonication (Branson Sonifier 250 equipped with a microtip: 1/8 "diameter, Danbury, Connecticut, USA, pulse: output 2, duty cycle: 35%) for 30 s.

A 3-day PEG broth preculture was used to inoculate 100 mL PEG broth with 1/75 volume of preculture. The biotransformations were started after 24 h incubation with the addition of 0.5 g L^−1^ of the substrates or glucose as a reference. The reactions were stopped after first products were detected (48 h for (*R*)-2-phenoxypropionic acid and 68 h for diclofenac (**1**)). The cells were then lysed as described above and the proteins of the samples analyzed and diluted as a whole cell suspension on a 10% and a 15% SDS-polyacrylamide gel.

The proteome analysis was carried out at the mass spectrometry service unit in Hohenheim (Germany, group of Dr. Pfannstiel) using an ACQUITY nano-UPLC system (Waters GmbH, Milford, USA) directly coupled to a LTQ-Orbitrap XL hybrid mass spectrometer (Thermo Fisher Scientific, Bremen, Germany) with following minor changes as described previously (29). Tryptic digests were separated on a 25 cm × 75 μm × 1.7 μm BEH 130 C18 reversed phase column (Waters GmbH) with gradient elution performed from 1% ACN to 50% ACN in 0.1% formic acid within 120 min. Identification of the proteins was based on a global NCBI database search applying the MASCOT search algorithm. The mass spectrometry proteomics data have been deposited to the ProteomeXchange Consortium via the PRIDE (30) partner repository with the dataset identifier PXD009664 and 10.6019/PXD009664.

### Preparative biotransformations

The filamentous fungi applied in this work were cultured in PEG broth, in which a better growth could be achieved in comparison to sabouraud medium. On PEG-agar, the hyphae were transferred once a month and incubated at 27°C for several days (*M. hiemalis*: 3 days, *B. bassiana*: 4 days, *C. nebularis*: 5 days). *B. bassiana* used in this work was modified by UV radiation to allow yeast-like growth in liquid culture (31). 50 mL of PEG broth was once inoculated with a loop tip, whereas *M. hiemalis* and *C. nebularis* were inoculated several times with hyphae to slightly disperse and therefore minimize lumpy mycelial growth. The fungi were cultured for 4 days at 27°C and 180 rpm and then transferred to 50 – 400 ml main cultures which were inoculated with 1/75 volume (*B. bassiana*) or with the filtered mycelia (*M. hiemalis* and *C. nebularis*) of the preculture. After 24 h of incubation, the biotransformations were started with 1 g L^−1^ substrate (50 g L^−1^ stock solution in DMSO) or in the case of diclofenac (**1**) with 0.6 g L^−1^ for 72 h or 144 h, respectively. At regular intervals, 500 μL samples were taken, centrifuged and the supernatant was used for analysis by GC-FID, HPLC-DAD or LC-MS. The biotransformations were carried out in biological duplicates starting from different agar plates.

### HPLC and LC-MS analytic

Diclofenac (**1**) and its metabolites were analyzed by HPLC and LC-MS as described elsewhere (21).

### GC analytic

Naproxen (**6**), ibuprofen (**8**) and mefenamic acid (**12**) were analyzed by gas chromatography. For this purpose, the samples (500 μL) were centrifuged and 250 μL of the supernatant initially acidified by 10 μL HCl prior extraction with the same volume of MTBE. The organic phase was then evaporated on a Genevac EZ-2 Plus Evaporator (Ipswich, UK) and resuspended in a mix of 50% MTBE and 50% BSTFA + TCMS (99: 1). The derivatization of the samples (150 μL) was carried out in GC vials at 70°C for 30 min. GC analysis was performed on a Shimadzu GC-2010 equipped with an AOC-20i autoinjector (Shimadzu, Nakagyo-ku, Japan). The samples were injected with a split of 20 (1 μL injection volume, injector temperature 250°C, carrier gas H2, 30 cm s^−1^) and separated by a DB-5 column (30 m × 0.25 mm × 0.25 μm, Agilent Technologies, Santa Clara, California, USA). The analytes were detected by a flame ionization detector (FID, detector temperature 330°C). For **6** and **12** the column temperature was maintained at 150°C for 1 min, increased to 280°C at a rate of 10°C min^−1^ and held for 1 min, raised to 320°C at a rate of 65°C min^−1^ and held for 3 min. For **8** the column temperature was maintained at 90°C for 1 min, increased to 280°C at a rate of 12°C min^−1^ and held for 1 min, raised to 320°C at a rate of 65°C min-1 and held for 3 min.

For initial product identification, the samples were run on GC-2010 GC-MS system equipped with Shimadzu GCMS-QP2010 detector and AOC-5000 autoinjector (Shimadzu, Nakagyo-ku, Japan) and helium as carrier gas (linear velocity 30 cm s^−1^). The analytes were separated on a DB-5 column and measured with identical temperature programs for GC-FID analysis. For the recording of the mass spectra, an ionization (EI, Electron Ionization) of 70 eV, an interface temperature of 250°C and an ion source temperature of 200°C was used. The detection of the mass fragments was finally carried out in the scan mode from 40 to 600 m/z.

### Product purification and identification

For product purification, the supernatant of the duplicates (2 × 50 mL or 2 × 400 mL) from preparative biotransformations were extracted twice with the identical volume of MTBE. In the case of biotransformations with substrate **6**, **8** and **12**, the supernatant was additionally acidified with HCl and afterwards extracted with MTBE. The organic phases were combined, evaporated to dryness and the residues were dissolved in 10 mL of acetonitrile and stored at 4°C for further use. The batches were finally purified by reverse phase semi-preparative HPLC on an Agilent 1200 System (St. Clara, California, USA) equipped with a G1311A quaternary pump, a HIP AS G1367B autosampler (1200 μL loop), a G1315D DAD detector and an analytical G1364C fraction collector. The separation of the analytes (injection volume: 1000 μL) was ensured using a Trentec Reprosil^®^ 100-5 C18 column (250 × 20 mm, 5 μm) from Dr. Maisch GmbH (Ammerbuch-Entringen, Germany). The mobile phases A and B were composed of water containing 0.1% formic acid and acetonitrile, respectively. Elution was done in a gradient mode at a flow rate of 5 mL min^−1^ and a column temperature of 20°C using following program: 85% A (0 min), 20% A (63 min), 20% A (67 min), 85% A (67.01 min), 85% A (83 min).

The elution of substrates and products were followed spectrophotometrically at a wavelength of 272 nm for **1**, 232 nm for **6**, 224 nm for **8** and 275 nm for **12**. Identical product fractions were combined, the solvent mixture of acetonitrile and water removed on a rotary evaporator and completely dried by a nitrogen gas flow. The remaining solid products were finally dissolved in *d*-methanol or *d*-DMSO and analyzed by ^1^H- and ^13^C-NMR to clarify the chemical structures. The purity of all samples were determined by GC or HPLC.

### NMR measurements

For the characterization of the purified products, NMR spectroscopy was used. ^1^H-,^1^H-COSY- and ^13^C-NMR spectra were recorded using a Bruker Avance 500 spectrometer at 500.15 and 125.76 MHz, respectively, or a Bruker Ascend 700 ™ spectrometer at 700.36 and 176.10 MHz respectively (both Bruker, Billerica, Massachusetts, USA). The chemical shift *δ* was measured in ppm (parts per million) and referred to TMS (tetramethylsilane) *δ* = 0 ppm as a standard. Unless otherwise stated, 10 mg of the product was used for NMR analysis. The NMR spectra are given in the supplemental material (Figures S2-S9).

5-Hydroxydiclofenac: ^1^H-NMR (500 MHz, CD3OD): *δ* (ppm) 3.71 (s, 2H; C-7), 6.37 (d, *J* = 8.8 Hz, 1H; C-3), 6.55 (dd, *J* = 8.8, 2.5 Hz, 1H; C-4), 6.72 (d, *J* = 2.5 Hz, 1H; C-6), 6.96 (t, *J* = 7.8 Hz, 1H; C-4’), 7.34 (d, *J* = 7.8 Hz 2H; C-3’ and 5’). ^13^C-NMR (126 MHz, CD3OD): *δ* (ppm) 39.2 (C7), 115.3 (ArCH-4), 118.3 (ArCH-6), 121.9 (ArCH-3), 124.1 (ArCH-4’), 129.2 (ArC-2), 129.3 (ArC-1’), 130.1 (2ArCH-3’ and 5’), 136.4 (ArC-1), 140.5 (2CCl), 154.1 (C-OH), 175.8 (COOH).

5-Hydroxyquinoneimine: ^1^H-NMR (500 MHz, (CD3)2SO): *δ* (ppm) 3.69 (s, 2H; C-7), 6.59 (dd, *J* = 10.1, 2.5 Hz, 1H; C-4), 6.74 (d, *J* = 10.1 Hz, 1H; C-3), 6.80 (d, *J* = 2.5 Hz, 1H; C-6), 7.25 (t, *J* = 8.2 Hz, 1H; C-4’), 7.58 (d, *J* = 7.6 Hz, 2H; C-3’ and 5’).

^13^C-NMR (126 MHz, (CD3)2SO): *δ* (ppm) 36.3 (C7), 123.1 (ArC), 126.6 (ArC), 128.6 (ArC), 129.1 (ArC), 133.1 (ArC), 133.5 (ArC), 143.4 (ArC-1’), 145.1 (ArC-1), 160.4 (ArC-2), 170.6 (COOH), 186.9 (C=O).

4’-Hydroxydiclofenac: ^1^H-NMR (500 MHz, CD3OD): *δ* (ppm) 3.7 (s, 2H, C-7), 6.26 (d, 1H, *J* = 8.0 Hz, C-3), 6.8 (t, 1H, *J* = 7.4, C-5), 6.88 (s, 2H, C-3` and C-5`), 7.02 (t, 1H, *J* = 7.5, C-4), 7.17 (d, 1H, *J* = 7.4, C-6).

^1^H-COSY (500 MHz, CD3OD): (7.17) *δ* (ppm) 3.7, 6.8, (7.02) *δ* (ppm) 6.26, 6.8.

^13^C-NMR (126 MHz, CD3OD): *δ* (ppm) 39.2 (C7‘), 116.4 (ArC), 116.9 (2ArC), 121.3 (ArC), 124.2 (ArC), 128.9 (ArC), 130.5 (ArC), 131.9 (2ArC), 133.7 (ArC), 145.5 (ArC), 156.3 (ArC-4`), 175.9 (COOH).

3’,4’-Dihydroxydiclofenac: ^1^H NMR (700 MHz, CD3OD): *δ* (ppm) 3.72 (s, 2H, C-7), 6.29 (d, 1H, *J* = 8.1 Hz, C-3), 6.80 (t, 1H, *J* = 7.6, C-5), 6.86 (s, 1H, C-5`), 7.02 (t, 1H, *J* = 7.9 Hz, C-4), 7.17 (d, 1H, 3.5 Hz, C-6).

6-*o*-Desmethylnaproxen: ^1^H-NMR (500 MHz, CD3OD): *δ* (ppm) 1.50 (d, 3H, *J* = 7.0 Hz, C-3), 3.81 (q, 1H, *J* = 7.2 Hz, C-2), 7.03 - 7.09 (m, 2H, C-8 and C-10) 7.35 (d, 1H, *J* = 8.5 Hz, C-11), 7.59 (d, 1H, *J* = 8.4 Hz, C-13), 7.64 (s, 1H, C-5), 7.67 (d, 1H, *J* = 8.8 Hz, C-6). ^13^C-NMR (126 MHz, CD3OD): *δ* (ppm) 15.4 (C7), 113.2 (ArC), 114.5 (ArC), 117.9 (ArC), 127.3 (ArC), 127.5 (ArC), 128.7 (ArC), 133.3 (ArC), 134.9 (ArC), 135.1 (ArC), 135.2 (ArC), 141.6 (ArC), 150.2 (ArC), 172.1 (COOH).

2-Hydroxyibuprofen: ^1^H-NMR (500 MHz, CD3OD): *δ* (ppm) 1.16 (s, 6H, C-12 and C-13), 1.43 (d, 3H, *J* = 7.2 Hz, C-3), 2.72 (s, 2H, C-10), 3.68 (q, 1H, *J* = 7.2 Hz, C-2), 7.18 (d, 2H, *J* = 8.0 Hz, C-5 and C-9), 7.22 (d, 2H, *J* = 7.9 Hz, C-6 and C-8).

^13^C-NMR (126 MHz, CD3OD): *δ* (ppm) 19.1 (C3), 29.2 (C12 and C13), 46.4 (C2), 50.3 (C10), 71.8 (C11), 128.0 (2x ArC), 131.9 (2 × ArC), 138.5 and 140.3 (C4 and C7), 178.6 (COOH).

1- Hydroxyibuprofen: ^1^H-NMR (700 MHz, (CD3)2SO): *δ* (ppm) 0.73 (d, 3H, *J* = 6.8 Hz, C-12) 0.85 (d, 3H, *J* =6.7 Hz, C-13), 1.32 (d, 3H, *J* = 7.1 Hz, C-3), 1.77 (oct, 1H, *J* = 16.4 Hz, C-11), 3.58 (q, 1H, *J* = 7.2 Hz, C-2), 4.19 (d, 1H, *J*= 6.2, C-10), 7.19 (d, 2H, *J* = 8.5 Hz, C-5 and C-9), 7.21 (d, 2H, *J* = 8.6 Hz, C-6 and C-8).

^13^C-NMR (176 MHz, (CD3)2SO): *δ* (ppm) 17.4 (C3), 18.2 and 18.5 (C12 and C13), 34.3 (C11), 44.4 (C2), 76.8 (C10), 125.9 and 126.1 (4 × ArC), 139.6 and 142.6 (C4 and C7), 175.3 (COOH).

1,2-Dihydroxyibuprofen: ^1^H-NMR (500 MHz, CD3OD): *δ* (ppm) 1.12 (s, 6H, C-12 and C-13), 1.43 (d, 3H, *J* = 3.4 Hz, C-3), 3.69 (q, 1H, *J* = 7.2 Hz, C-2), 4.43 (s, 1H, C-10), 7.26 (d, 2H, *J* = 7.8 Hz, C-5 and C-9), 7.34 (d, 2H, *J* = 8.0 Hz, C-6 and C-8).

^13^C-NMR (126 MHz, CD3OD): *δ* (ppm) 19.1 (C3), 25.3 and 26.0 (C12 and C13), 46.7 (C2), 74.0 (C11), 81.6 (C10), 127.8 (2 × ArC), 129.2 (2 × ArC), 141.7 and 141.8 (C4 and C7). COOH group is barely visible.

3’-Hydroxymethylmefenamic acid: ^1^H-NMR (500 MHz, (CD3)2SO): *δ* (ppm) 2.11 (s, 3H, C-8’), 4.53 (s, 2H, C-7’), 6.68 – 6.73 (m, 2H, C-2 and C-4), 7.18 – 7.25 (m, 3H, C-4’, 5’ and 6’), 7.31 (t, 1H, *J* = 7.5 Hz, C-3), 7.89 (d, 1H, *J* = 7.5 Hz, C-5).

^13^C-NMR (126 MHz, (CD3)2SO): *δ* (ppm) 12.4 (C8‘), 61.3 (C7`), 111.6 (ArC), 112.9 (ArC), 116.3 (2ArC-3 and 4‘), 122.6 (ArC), 123.5 (ArC), 125.9 (ArC), 129.9 (ArC), 131.6 (ArC), 134.0 (ArC), 138.2 (ArC), 141.8 (ArC), 170.0 (COOH).

3’-Carboxymefenamic acid: ^1^H-NMR (500 MHz, (CD3OD): *δ* (ppm) 2.43 (s, 3H, C-8’), 6.71 (t, 1H, *J* =7.5 Hz, C-4), 6.75 (d, 1H, *J* = 4.3 Hz, C-2), 7.28 (q, 2H, *J* = 7.3 Hz, C-3, and C-5’), 7.46 (d, 1H, *J* = 7.8 Hz, C-4’), 7.62 (d, 1H, *J* = 7.8 Hz, C-6’), 7.98 (d, 1H, *J* = 8.0 Hz, C-5).

^13^C-NMR (126 MHz, (CD3OD): *δ* (ppm) 15.4 (C8‘), 113.2 (ArC), 114.5 (ArC), 117.9 (ArC), 127.3 (ArC), 127.5 (ArC), 128.7 (ArC), 133.3 (ArC), 134.9 (ArC), 135.1 (ArC), 135.2 (ArC), 141.6 (ArC), 150.2 (ArC), 172.1 (2 × COOH).

## Results and Discussion

### Identification of microbial biocatalysts for NSAID oxyfunctionalizations

In addition to the already published organism *Beauveria bassiana*, which is able to perform one of the most effective hydroxylation processes in biocatalysis, the production of (*R*)-2- (4-hydroxyphenoxy)propionic acid and a variety of other hydroxylation reactions (31, 32), we identified two other filamentous fungi as further novel NSAID accepting biocatalysts. One of these strains was isolated from soil samples close to Ludwigshafen (Germany) during previous diclofenac (**1**) degradation experiments (33). The third filamentous fungus was isolated by coincident as a contamination strain out of a 5-hydroxydiclofenac (**3**) standard, diluted in M9 minimal medium, after 1.5 years of aerobic incubation. Both fungi were assigned by the universal biomarkers LSU (28S nuclear ribosomal large subunit rRNA gene) and ITS (internal transcribed spacer). Sequence matching with the fungal barcoding and the NCBI database revealed that one strain was > 99% consistent with the S1 organism *Clitocybe nebularis* (confirmed by LSU and ITS biomarkers). For the other organism, the LSU biomarker found a > 98% similarity with the S1 organism *Mucor hiemalis*.

### Biotransformations with diclofenac

The fungi were cultivated in 400 mL PGE broth and the biotransformations were done in duplicates with a starting concentration of 0.6 g L^−1^ diclofenac (**1**). The reaction was monitored over time by HPLC-DAD and stopped after 144 h (Figure 1). All products were purified by semi-preparative HPLC for further detailed structure identification by ^1^H and ^13^C NMR. Similar to the distribution of products in humans, the 4’-position (**2**) was favoured in the biotransformations with the fungi, which was also observed with P450s and microorganisms in previous studies (Scheme 1, Table 1). In comparison to *C. nebularis* which is able to catalyze a mixture of the metabolites 4’-hydroxydiclofenac (**2**, 60%), 5-hydroxydiclofenac (**3**, 30%) and the corresponding quinoneimine (**4**, 10%) with 50.6% isolated yield, *M. hiemalis* selectively oxyfunctionalized diclofenac (**1**) yielding 61.7% of 4’-hydroxydiclofenac (**2**) after purification. High isolated yields of 55.3% were additionally achieved using *B. bassiana*. Next to the major product 4’-hydroxydiclofenac (**2**, 90%), a 3’,4’-dihydroxydiclofenac (**5**) metabolite was identified that has not been characterized and published yet. In studies concerning the fungi *Epicoccum nigrum* and *Cunninghamella elegans*, preparative biotransformations were performed resulting in product titers of 16 and 57 mg L^−1^, respectively (34, 35). In contrast, the filamentous fungi identified in the present work were able to obtain significantly higher values comparable with biotransformations using P450 RhF and P450 BM3 (Table 1).

**FIG 1.**
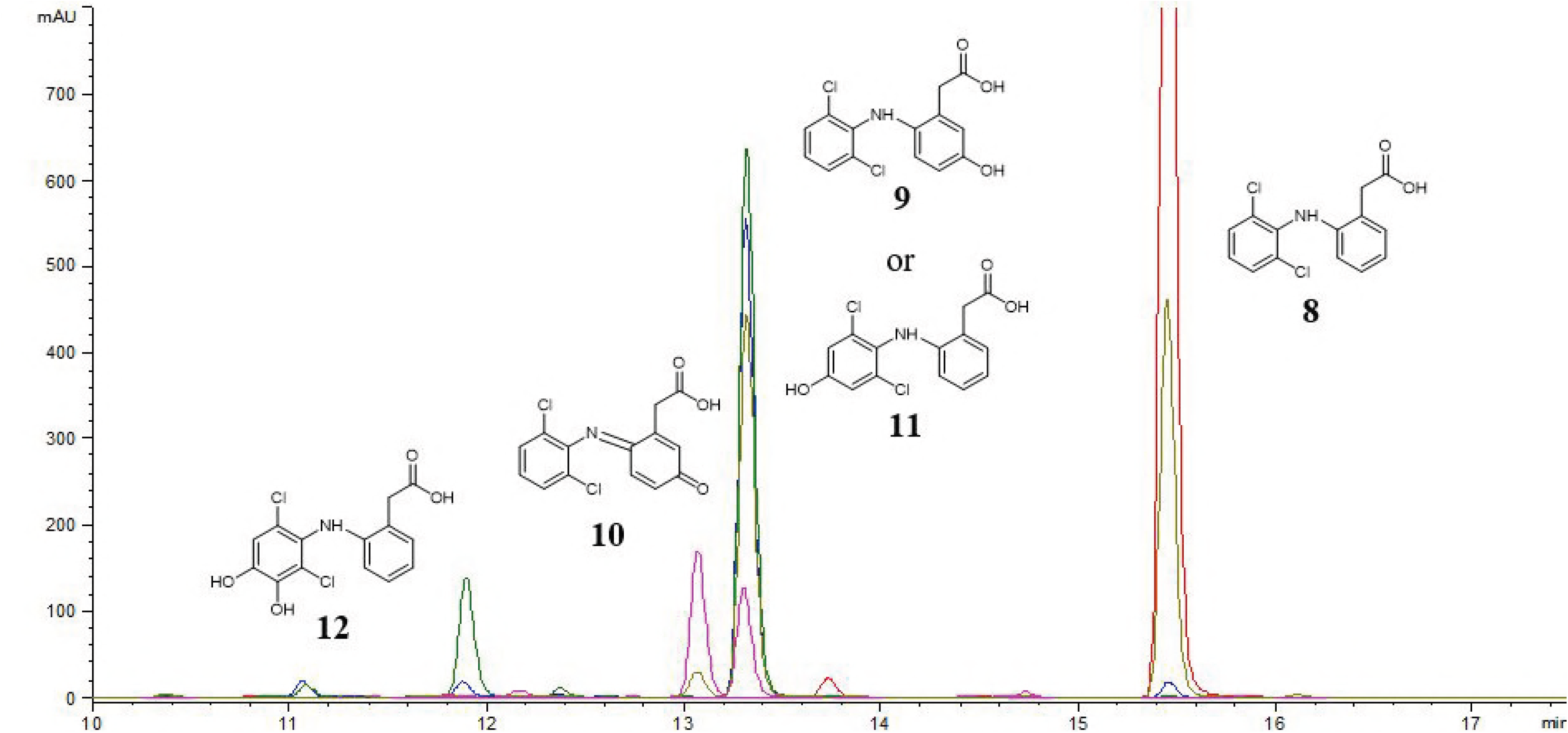
Diclofenac (**1**) biotransformations with the filamentous fungi *Beauveria bassiana*, *Clitocybe nebularis* and *Mucor hiemalis*. The reactions were carried out in 400 mL of PEG broth with 0.6 g L^−1^ substrate for 144 h at 27°C and 180 rpm. **Red**: negative control without cells; **Magenta**: standard of 4’-hydroxydiclofenac (**2**), 5-hydroxydiclofenac (**3**) and 5-hydroxyquinoneimine (**4**); **Green**: *Beauveria bassiana*; **Blue**: *Mucor hiemalis*; **Ocher**: *Clitocybe nebularis*. The retention times for the different metabolites are: 11.9 min (**5**); 13.1 min (**4**); 13.3 min (**2** and **3**); 15.5 min (**1**).

**SCHEME 1.**
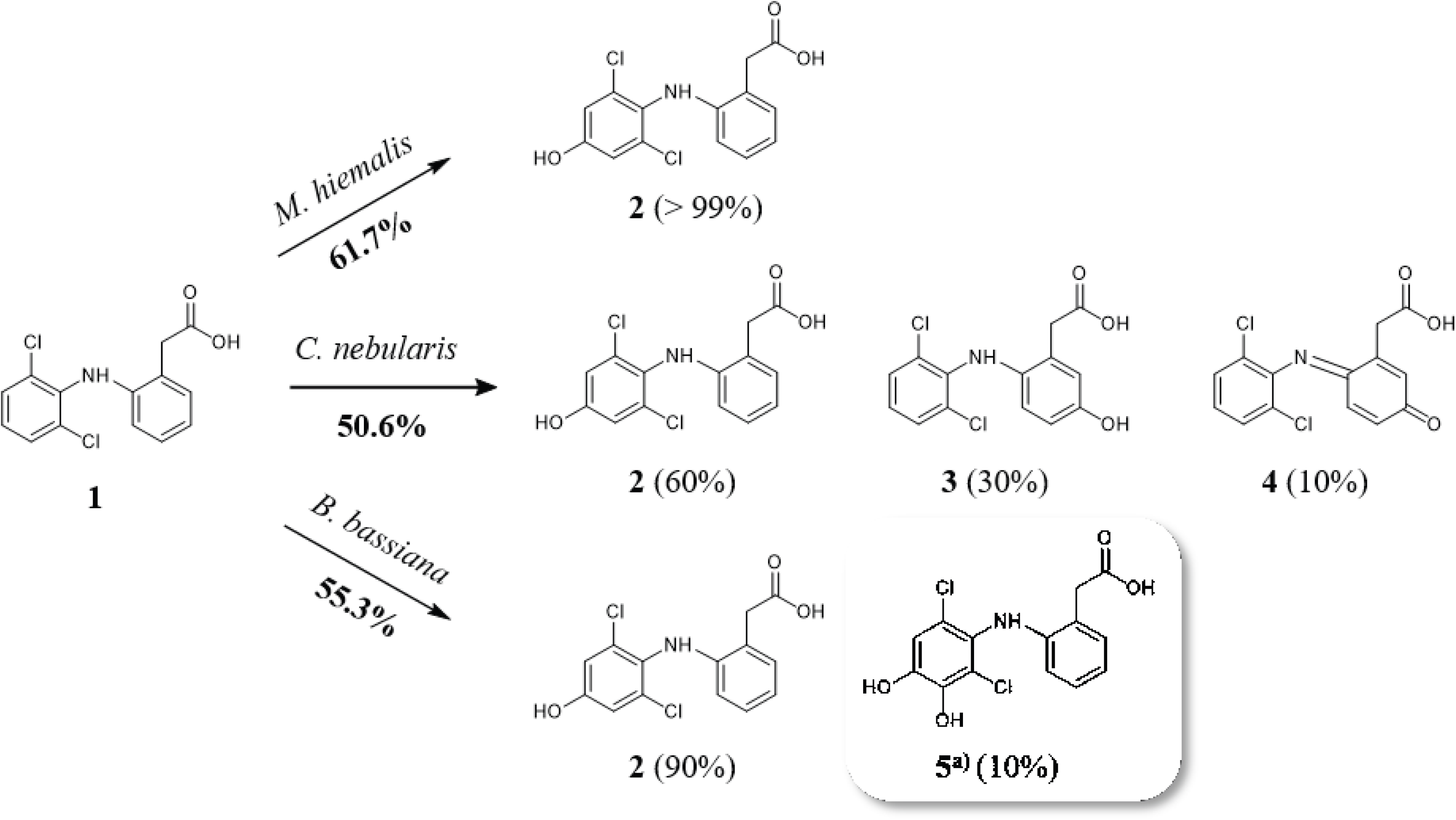
Diclofenac metabolites (**2** – **5**) synthesized by different filamentous fungi as biocatalyst using 0.6 g L^−1^ substrate. The isolated yields after extraction and purification *via* semi-preparative HPLC and the product distribution in brackets are given. The metabolites 4’-hydroxydiclofenac (**2**), 5-hydroxydiclofenac (**3**), 5-hydroxyquinoneimine (**4**), 3’,4’-dihydroxydiclofenac (**5**) were identified by NMR. ^a)^Not yet observed diclofenac metabolite.

**TAB 1.**
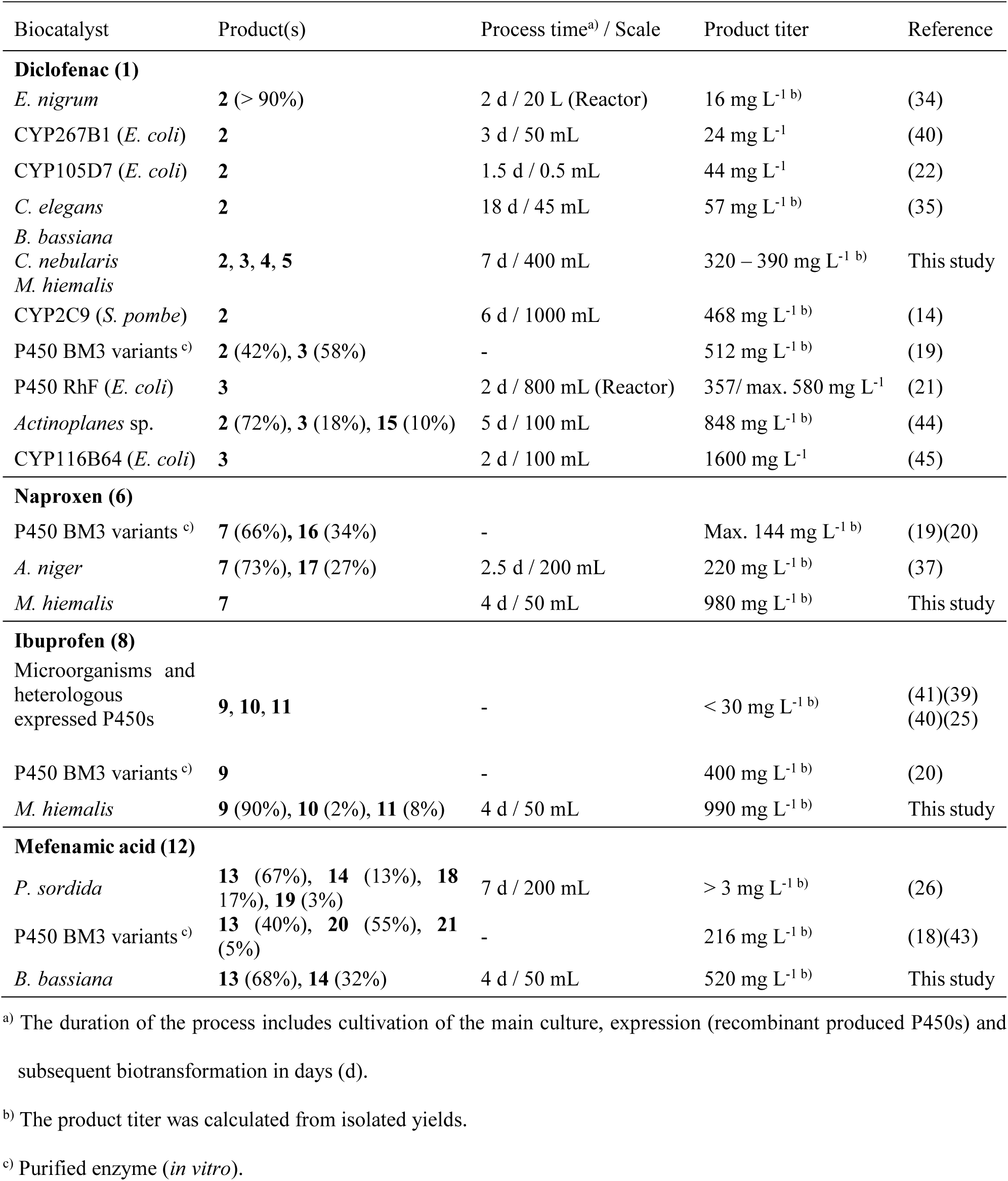
Summary of biotechnological processes to produce NSAID metabolites. 4’-hydroxydiclofenac (**2**); 5-hydroxydiclofenac (**3**); 5-hydroxyquinoneimine (**4**); 3’,4’-dihydroxydiclofenac (**5**); 4’,5-dihydroxydiclofenac (**15**); 6-*o*-desmethylnaproxen (**7**); 2- hydroxyibuprofen (**9**); 1-hydroxyibuprofen (**10**); 1,2-dihydroxyibuprofen (**11**); 3’-hydroxymethylmefenamic acid (**13**); 3’-carboxymefenamic acid (**14**); 2-Acetyl-6- methoxynaproxen (**16**); 7-hydroxynaproxen (**17**); 3’,5-dihydroxymefenamic acid (**18**); 3’,6’-dihydroxymefenamic acid (**19**); 4’-hydroxymefenamic acid (**20**); 5-hydroxymefenamic acid (**21**).

### Proteome analysis of B. bassiana

A common question that is often raised as a negative element for the use of microbial cell factories is which enzymes are involved in the detected biotransformation. In this respect, we aimed to identify 450s, which are responsible for the oxyfunctionalization of diclofenac (**1**) and (*R*)-2-phenoxypropionic acid in *B. bassiana* using omics studies. As a proof of concept, we analyzed biotransformation samples whether we can identify differences by a proteome analysis depending on the substrate. Therefore, samples of biotransformations with the corresponding substrates and glucose as negative control were processed to obtain the organisms proteome. By applying the proteome analysis clear evidence of P450s (implied by the number of detected and assigned fragments) which were upregulated with the specific substrates, were achieved drastically reducing the high number of 83 putative P450s in *B. bassiana* (36). With diclofenac (**1**) as substrate two P450s, CYP548A5 and CYP51F1, were exclusively expressed compared to the samples exposed with the other substrates (Table 2). In case of (*R*)-2-phenoxypropionic acid as substrate, CYP52A10 seems to be strongly upregulated while CYP684A2 was induced exclusively. In eukaryotes, only one bifunctional NADPH-reductase serves as electron delivering partner that was also found in all samples. In this work, the proteome analysis has thus proved to be a targeted method for identifying involved proteins in eukaryotes that can subsequently lead to valuable new heterologous expressed biocatalysts.

**TAB 2.** Results of the proteome analysis of the biotransformations with *Beauveria bassiana* carried out with different substrates. The numbers stand for the quantity of clearly attributable peptide fragments to a protein. A higher number contributes to a higher probability that the protein is actually present in the sample. A sample with glucose as substrate served as a reference.

|  | Glucose | Diclofenac (**1**) | (*R*)-2-phenoxypropionic<br>acid |
| --- | --- | --- | --- |
| NADPH-CYP-Reductase | **14** | **9** | **10** |
| CYP52A10 | **2** | 0 | **11** |
| CYP52A13 | **5** | 0 | 0 |
| CYP548A5 | 0 | **5** | 0 |
| CYP51F1 | 0 | **1** | 0 |
| CYP684A2 | 0 | 0 | **2** |

### NSAID biotransformations applying the identified microbial biocatalysts

To investigate the potential of the microbial biocatalysts further we increased the NSAID substrate scope of the fungi strains to oxyfunctionalize naproxen (**6**), ibuprofen (**8**) and mefenamic acid (**12**). In some cases, high titers and full conversion towards the investigated NSAIDs were achieved. For example, *M. hiemalis* demethylated naproxen (**6**) to the human metabolite 6-*o*-desmethylnaproxen (**7**) with an isolated yield of over 99% using 1 g L^−1^ of substrate (Scheme 2). In comparison the *in vitro* biotransformations with P450 BM3 variants and the strain *A. niger* showed much lower selectivity and isolated yield (Table 1) (19, 20, 37).

**SCHEME 2.**
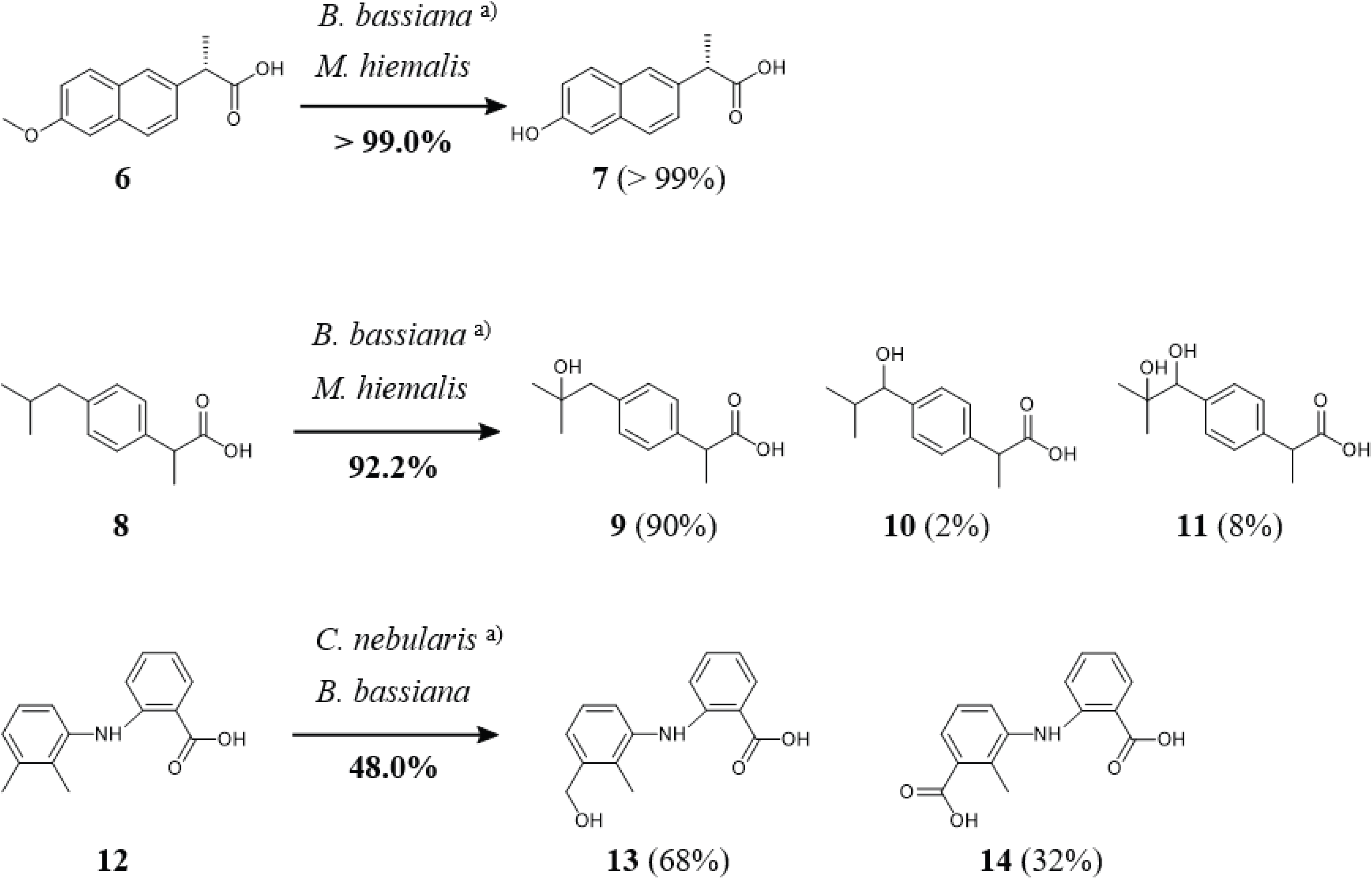
NSAID-metabolites synthesized by different filamentous fungi as biocatalyst using 1 g L^−1^ substrate. The isolated yields after extraction and purification *via* semi-preparative HPLC and the product distribution in brackets are given. The metabolites 6-*o*-desmethylnaproxen (**7**), 2-hydroxyibuprofen (**9**), 1-hydroxyibuprofen (**10**), 1,2-dihydroxyibuprofen (**11**), 3’-hydroxymethylmefenamic acid (**13**) and 3’-carboxymefenamic acid (**14**) were identified by NMR. ^a)^Products were not purified; hence no isolated yield and exact products are given.

Furthermore, *M. hiemalis* almost completely hydroxylated 1 g L^−1^ ibuprofen (**8**) to the metabolites 2-hydroxyibuprofen (**9**, 90%) 1-hydroxyibuprofen (**10**, 2%), and the double hydroxylated secondary product 1,2-dihydroxyibuprofen (**11**, 8%) (Scheme 2). The highest product titer so far was achieved by a P450 BM3 variant *in vitro* with 0.4 g L^−1^ of 2-hydroxyibuprofen (**9**) (Table 1) (17, 20, 38). Microorganisms such as *Trametes versicolor* and *Nigospora sphaerica* or the P450s CYP2C9, CYP267A1 and B1 had significantly lower product yields (> 30 g L^−1^) (25, 39–41). Using the anthranilic acid derivative mefenamic acid (**12**) as substrate, the highest activity was achieved by *B. bassiana* with an isolated yield of 48% at 1 g L^−1^ substrate concentration (Scheme 2). Next to the 3’-hydroxymethylmefenamic acid (**13**) as main product (68%), the product 3’-carboxymefenamic acid (**14**) was catalyzed. In humans, **12** is mainly converted to **13** by CYP2C9 which undergoes further oxidation to the carboxy metabolite **14** (42). An evolved P450 BM3 variant generated besides 3’-hydroxymethylmefenamic acid (**13**) the other human P450 catalyzed metabolites 4’- and 5-hydroxymefenamic acid (**20** and **21**) which are hydroxylated at the aromatic rings (18, 43). In this case, *in vitro* product titers of 216 mg L^−1^ were below those obtained with *B. bassiana* (Table 1). In degradation experiments, the wild-type organism *Phaenerochaete sordida* (white-rot fungus) also produced different mefenamic acid metabolites with significantly lower yields of less than 3 mg L^−1^ (26).

Overall, the filamentous fungi of the present study achieved high product titers in the biotransformations with the NSAIDs. Applying a semi-preparative HPLC purification method, all NSAID metabolites could be isolated in high yields and high purities. Moreover, reproducible product titers were obtained at 50 mL or 400 mL scale exhibiting the possibility to further increase the yields by improving the cultivation and biotransformation procedure. One drawback of filamentous fungi as catalyst for biotransformations is their mycelial growth. This particular cell shape minimizes the possible surface to the medium and thus the mass transfer of substrate solubilized in the medium and the cells. Especially cells located inside the mycelia may therefore be unable to participate in biocatalysis, but against our expectation, the clump-forming organism *M. hiemalis* has achieved the highest yields so far.

Compared to already published biotechnological processes the total process time with the filamentous fungi of 4 days or 7 days for diclofenac (**1**), respectively, were in accordance with most of the studies (Table 1 and 2). However, a simple cultivation and biotransformation setup using fungi with less steps are main benefits, besides the achieved high product yields. In detail, compared to P450-based reactions, less requirements on the process were necessary since issues associated with an often complex P450 system like biocatalyst expression, stability or cofactor dependency are negligible.

In conclusion, the results of the present work demonstrate the potential of microbial biocatalysts for NSAID oxyfunctionalizations. Often, low activity, difficult handling, or high by-product formation are illustrated as criteria against the use of microorganisms (40). In contrast, the microorganisms identified in this work are easy to cultivate, inexpensive and are able to produce high valued metabolites in promising yields. The applied and identified filamentous fungi serve with their versatile metabolism as an interesting platform for novel cell factories to produce valuable NSAID metabolites in an effective and cheap way or can help to develop new strategies in drug design.

## Acknowledgements

The research leading to these results has received funding from the European Union’s Seventh Framework Programme for research, technological development and demonstration under grant agreements no 613849 (BIOOX). We also gratefully acknowledge financial support from the Landesgraduiertenförderung (LSFG) Baden-Württemberg. We thank Berit Würtz and Dr. Jens Pfannstiel for conduction and helpful advice concerning the proteome analysis.

## Abbreviations

GC: gas chromatography
HPLC: high-performance liquid chromatography
MS: mass spectrometry
NMR: nuclear magnetic resonance
NSAID: non-steroidal anti-inflammatory drug
PEG: potato-extract-glucose
P450: cytochrome P450 monooxygenase
P450 BM3: cytochrome P450 from *Bacillus megaterium*
P450 RhF: cytochrome P450 from *Rhodococcus* sp. strain NCIMB 9784

